# Cooperative efforts on developing vaccines and therapies for COVID-19 Cooperative efforts for COVID-19

**DOI:** 10.1101/2020.09.06.282145

**Authors:** Fernanda Gisele Basso, Alex Fabianne de Paulo, Geciane Silveira Porto, Cristiano Gonçalves Pereira

## Abstract

Health organizations have always sought partnership to join competencies in innovation, even with fierce competition in this sector. In this pandemic moment it is relevant to observe how organizations behave to seek quick and safe answers. The present research analyzes how the cooperation networks were set off considering the clinical trials on therapies and vaccines that were developed specifically to treat or prevent COVID-19. Social Network Analysis technique was used to build cooperation networks and apply metrics that characterize these connections. There was an evaluation of statistics of Strength of cooperation and Unilateral dependence of cooperation that identify the cooperation strength between two organizations, and the dependence of this relations. A total of 415 clinical trial were identified, of which 42% are in cooperation. From organizations that have partnership, firms are the first, followed by universities. We extracted the main categories that concentrate 74% of partnerships in the trials of antibody, and vaccine. Several organizations cooperate in multiple categories of trials, evidencing the efforts to focus on different strategies to treat the disease. We found high strength of cooperation and an assimetryc dependency between partners, which can be assigned to specialized models of partnership and it occurs in competitive enviroments like this pandemic moment. Cooperation were not limited to geographical proximity and the advent of Chinese players can represent a new change in the biotechnological development axis. Finally, the challenge of finding therapeutic or immunological solutions for COVID-19 demonstrates a clear composition of cooperation groups that complement their skills to manage organizational strategies to beat the pandemic. In this new paradigm, there can be partnerships not only in clinical trial but also in pre-competitive technologies development. This experience is expected to change the way of organizations define their R&D strategies and start to adopt more a collaborative innovation model.

## Introduction

The estabilishment of the partnership between firms and Institutes of Science and Technology (IST) has brought significant contributions to the transformation of knowledge resulting from technological development. The use of knowledge generated in universities by the productive sector are the driver for the development of new technologies whose transfer consists of a complementary path to reach a higher technological level [1]. From the perspective of Open Innovation (OI), organizations improve their capabilities by combining the internalization and ourtsourcing of resources [2] and establishing partnerships with other firms, customers, suppliers, or IST, which occur at different levels of affinity and complexity. The level of maturity that can be achieved in OI is the strong prioritization of the development of partnerships [3].

Technological cooperation networks, which are formed by groups of heterogeneous organizations [4], can be understood from the perspective of their structure: a horizontal pattern when partners from different sectors collaborate or a vertical pattern when the actors are from the same production chain [5-6]. Thus, organizations are grouped based on the formation of partnerships with firms, government agencies, investors, industrial associations, and research institutions [7]. Organizations which are in cooperation networks that are more heterogeneous tend to be associated with better economic performance [8]. In addition, highly clustered communities induce the feeling of trust [9], which is relevant when a partnership is associated with discoveries commercially attractive and with high profit expectations.

Thus, selecting of the best partners is one of the main challenges for the success of businesses in cooperation [10]. A deep understanding of the cooperation networks can be important and can provide insights of the pattern of relationships affecting the innovation performance of their members; however, the overflow of knowledge may not be equally accessible by or appropriate for all members [11].

Thus, regarding the development of drugs for COVID-19, the cooperation networks resulting from clinical trial (CT) may emerge as an essential element, allowing organizations to share their results and fulfill the expectations of the drugs efficacy in a more intensive and rapid manner. The decision on which partner to choose to develop an R&D&I project in this sector are frequently drived by complimentray competences to beat the pandemic faster but also focusing on firms’ strategies to optimizing their CT portfolios. Identifying the most promising technology in such a short time compared to traditional regulatory parameters, then scaling them up and distributing these therapies and/or vaccines are a major challenge for investors and firms, regulatory agencies, and academics. In this context, cooperation networks during the pandemic can be perceived as performance indicators when joining capabilities are enforced to fight the disease and provide a solution for the pandemic.

Efforts to seek partnerships and create new alliances to fight COVID-19 have brought numerous challenges. There are risks due to the need to redirect the efforts of research teams despite working on the ongoing funding (i.e. for other diseases). There may be delays, discontinuities or even difficulties in maintaining adherence to original protocols, which impairs statistical consistency [12]. Notably, the pressure to obtain rapid results cannot lead to disregard the safety and integrity of the trials; therefore, the organizations involved are seeking maximum effectiveness in the shortest possible time by incorporating knowledge accumulated during other pandemics [13-14]. In addition, concern regarding the accurate monitoring of numerous CTs while avoiding redundancies has led to the creation, via artificial intelligence, of a real-time dashboard for COVID-19 CTs [15].

Given the extent and severity of COVID-19, numerous studies under different perspectives have sought to analyse developments related to the pandemic. However, the dynamics of cooperation, which can accelerate the development of a vaccine or drug that is effective against this disease, have not yet been explored. This study seeks to deply understand the cooperation networks of organizations that have combined efforts to create and test new drugs to fight COVID-19, based on the 415 CT mapped until July 2020.

With this purpose, social network analysis (SNA) were used to map the interactions between organizations, understand the determinants of cooperation and to allows the analysis of a significant sample of CT resulting from collaborative development [16-17]. Notably, within the set of studies on cooperation networks, the adoption of SNA using CT data is a strategy little explored in the literature. Therefore, this study contributes to understand the profile of entities that cooperate the most in a pandemic status and to highlight which technologies are more prone to the need of join forces to complete the clinical and the technology development. During the pandemic, the power of science and technology are in the spotlight, and the world needs a quick and effective solution to beat COVID-19. In this regard, our study demonstrate that the estabilishemet of partnerships and the networks made by that shows the efforts of the firms to developed highly complex drugs overcoming the barrier of competition loosen the burocracy embedded in the agreements and speeding up the start of clinical development, all in favor of the health of the population.

## METHOD

This study uses SNA as a method to identify and build cooperation networks for CTs focused on the development of drugs and vaccines for COVID-19. The operational definition of the main constructs used for innovation [4], SNA [18], and clinical trials [19] are detailed in the supporting material (S1 Appendix). The study used data sources from Bio Century, clinicaltrials.gov, and Policy Cures Research collected until July 10th, 2020.

The organizations were classified into firms, universities, hospitals, research institutes (RIs) and government. The categories of clinical trials were grouped into eight categories (S1 Table). The sponsors, which are organizations that provide funding, were excluded from the networks because their participation was assumed to occur only through funding and not in the technological development and trials themselves.

For the construction of cooperation networks, it was assumed that organizations signed agreements and/or treaties to develop specific studies, establishing joint ownership of the results and the new drug or vaccine, with the purpose of forming an alliance for innovation [20]. CTs with 2 or more organizations were label standardized and analysed using Gephi for graphical representation and calculation of the network metrics. The connections (edges) refer to the (non-directional) relationship of partnerships in each CT.

The Strength of cooperation (SC) was adopted to identify the affinity of the cooperation relationship between two organizations, and the unilateral dependence on a cooperation relationship (UDC) was used to verify the asymmetry collaboration relationship [21]. SC and UDC values vary between 0 and 1, and the results of each relationship are distributed in quartiles (Q) as follows: Q1 = 0 to 0.25; Q2 = 0.26 to 0.5; Q3 = 0.51 to 0.75, and Q4 = 0.76 to 1. To avoid bias in the analysis, one-to-one relationships were suppressed because the metric values would automatically have a maximum value (S2 Appendix).

## RESULTS

One of the main facilitators and drivers for the technology and clinical development are the funding agencies globally. Regarding strategic funding activity, five main sponsors stand out: the Biomedical Advanced Research and Development Authority (BARDA), the Coalition for Epidemic Preparedness Innovations (CEPI), the Bill & Melinda Gates Foundation, the Bloomberg Foundation and the National Health Service (NHS), which invested approximately two billion dollars (Fig 1).

**Fig 1.**
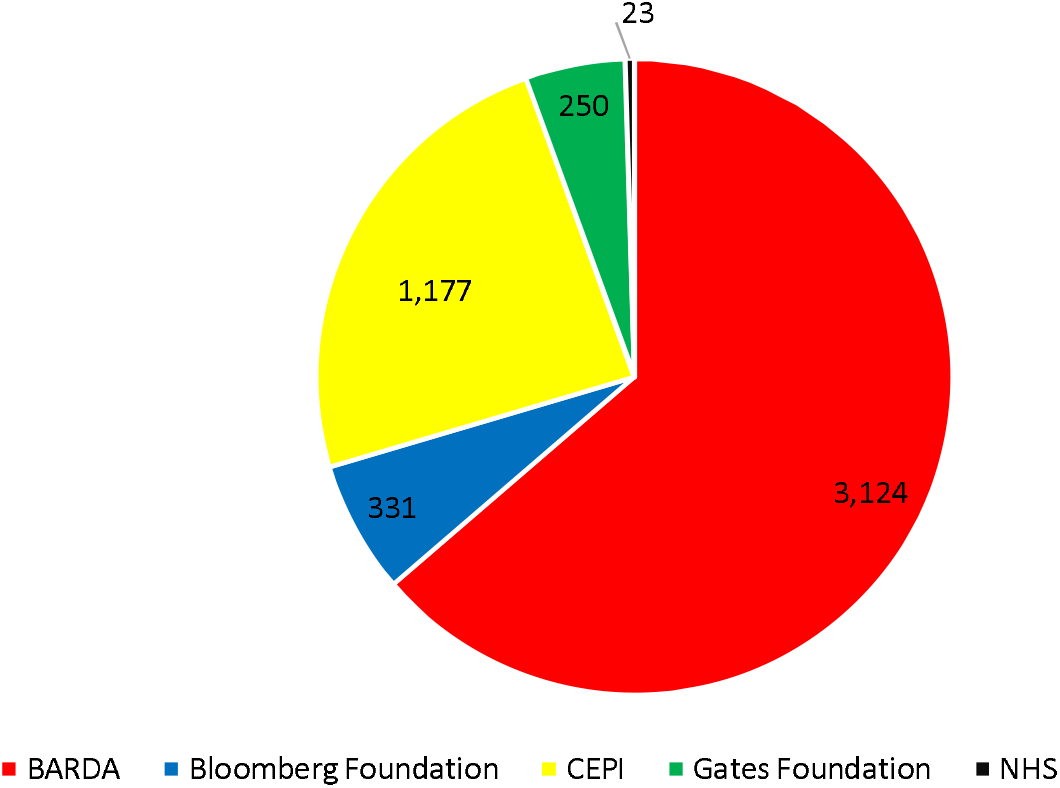
Main sponsor for funding COVID-19 CT in millions of dollars.

### Clinical trials involving vaccines and new therapies for COVID-19

A total of 415 ongoing CTs involving drugs and vaccines for COVID-19 were identified, of which 42% are in cooperation and 58% are being tested by only one organization (without cooperation). The most tested therapeutic categories involve antibodies, vaccines, and proteins, which represent 89% of the total CTs in progress. The following categories stand out based on cooperation relevance (Fig 2), in decreasing order: siRNA (73%), vaccines (47%), protein-based technologies (45%), nucleic acid-based technologies (43%), antibodies (41%), and cell therapy (36%). Conversely, despite technological effort, only 7% of trials involving small molecules have cooperation on CT (S2 Table).

**Fig 2.**
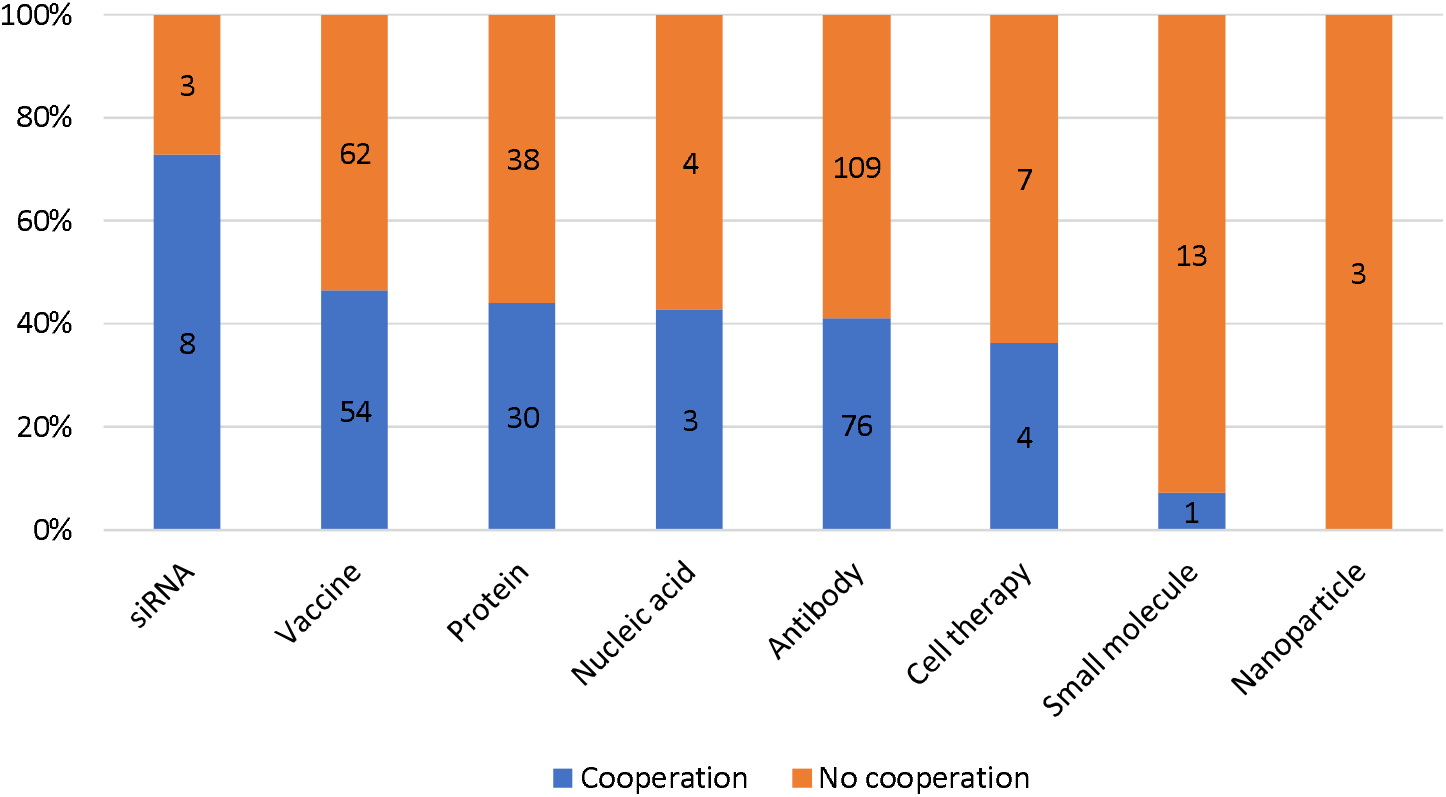
CT categories executed with or without cooperation.

Notably, organizations seek to accelerate technological development by establishing partnerships [22]. In the case of COVID-19, time is crucial because obtaining effective results can generate a broad competitive advantage in this market, in addition to contributing to the normal resumption of post-pandemic activities. Regarding the distribution of the types of organizations that cooperate by category (Fig 3), a greater diversity of partnerships was observed in antibodies, followed by vaccines and proteins, because these categories address complex therapies and require complementarity among several disciplines (e.g., adjuvants in vaccines and expression systems for proteins). The firms are involve in all categories, but with higher proportions on antibodies and exclusively in siRNA. It is important to highlight the effort of hospitals regarding antibody therapies, cooperating in vaccines candidates, protein-based and nucleic acid-based tehrapies in clinical trials. Universities and RIs are also relevant and are present in collaborations for most vaccine categories. The government acts only in clinical trials related to antibody-based therapies. Hospitals stand out because they are fundamental for the development of CTs as working as R&D and clinical centers. Another important aspect is that 62% of the firms (191) opted for the development of CTs in cooperation rather than alone, which may be associated with the need for quick responses arising from the complementarity of technological skills obtained through the establishment of partnerships.

**Fig 3.**
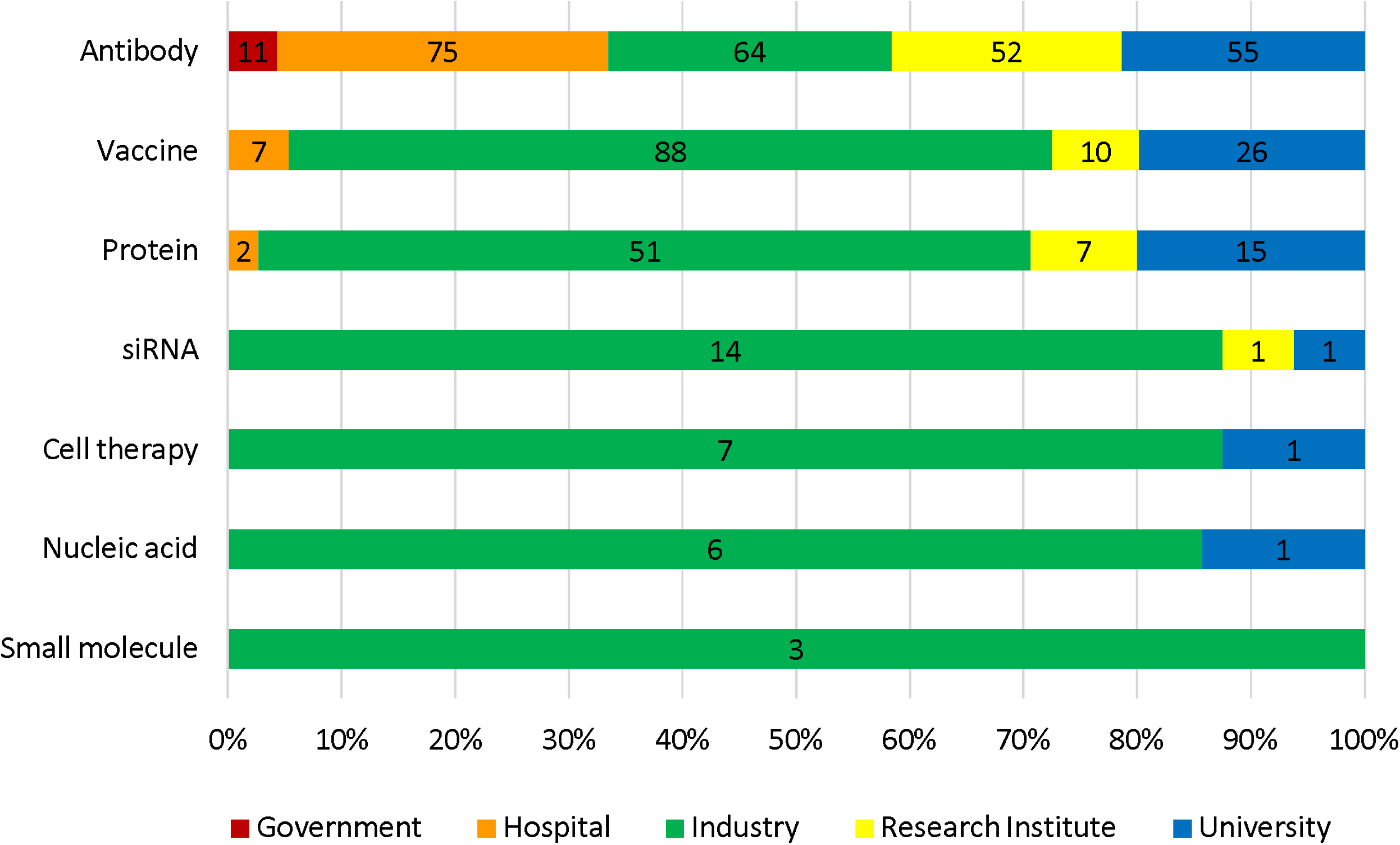
CT cooperation by categories and organizations type.

Because clinical adoption and commercial success are due to the incorporation degree of existing practices in innovation processes [23], the diffusion of disruptive technologies in this field may encounter greater challenges. Thus, there may be some difficulties in adhering to technologies involving siRNA because there are few firms with expertise in their development which delays the approval of drugs. It is difficult to correctly deliver siRNA treatments without the drugs being degraded by nuclease enzymes and without promoting side effects [24]. This technological challenge is also observed in treatments involving cell therapy [25].

### Cooperation analysis

The general cooperation network is constituted by 177 CTs (Fig 4), and 407 organizations establish 701 cooperation relationships (S3 Table), indicating specific partnerships. There are exception for 12 collaboration involved in two CTs and two partnerships involved in three CTs.

**Fig 4.**
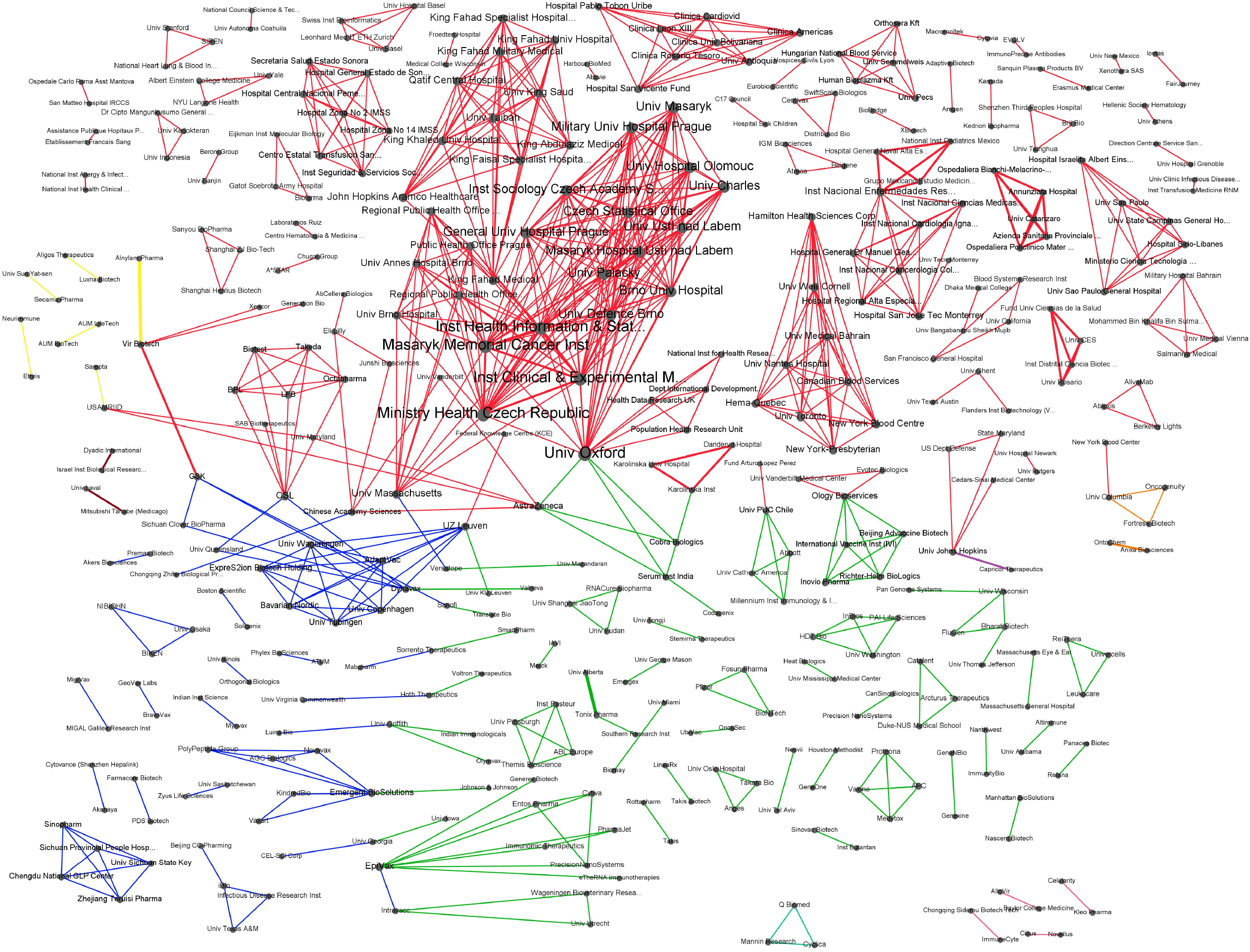
General cooperation network for COVID-19 therapies and vaccines. Edges represent CT categories. Legend: Red = Antibody; Dark blue = Protein; Dark green = Vaccine; Grey = Nucleic Acid; Orange = Small molecule; Dark red = Cell Therapy; Yellow = SiRNA; Light green = Protein/Antibody; Light blue = Protein/Vaccine

By dividing the network by technological categories (Fig 5), 58 connected groups related to technologies involving antibodies were identified. Most of them are hospitals that work together with several organizations, including other hospitals, in the same convalescent plasma CT. In general, these cooperation relationships were carried out mainly by geographic proximity and involving government agencies. Despite increasing globalization, regional action is relevant to innovation networks because it facilitates the exchange of knowledge between organizations [26].

**Fig 5.**
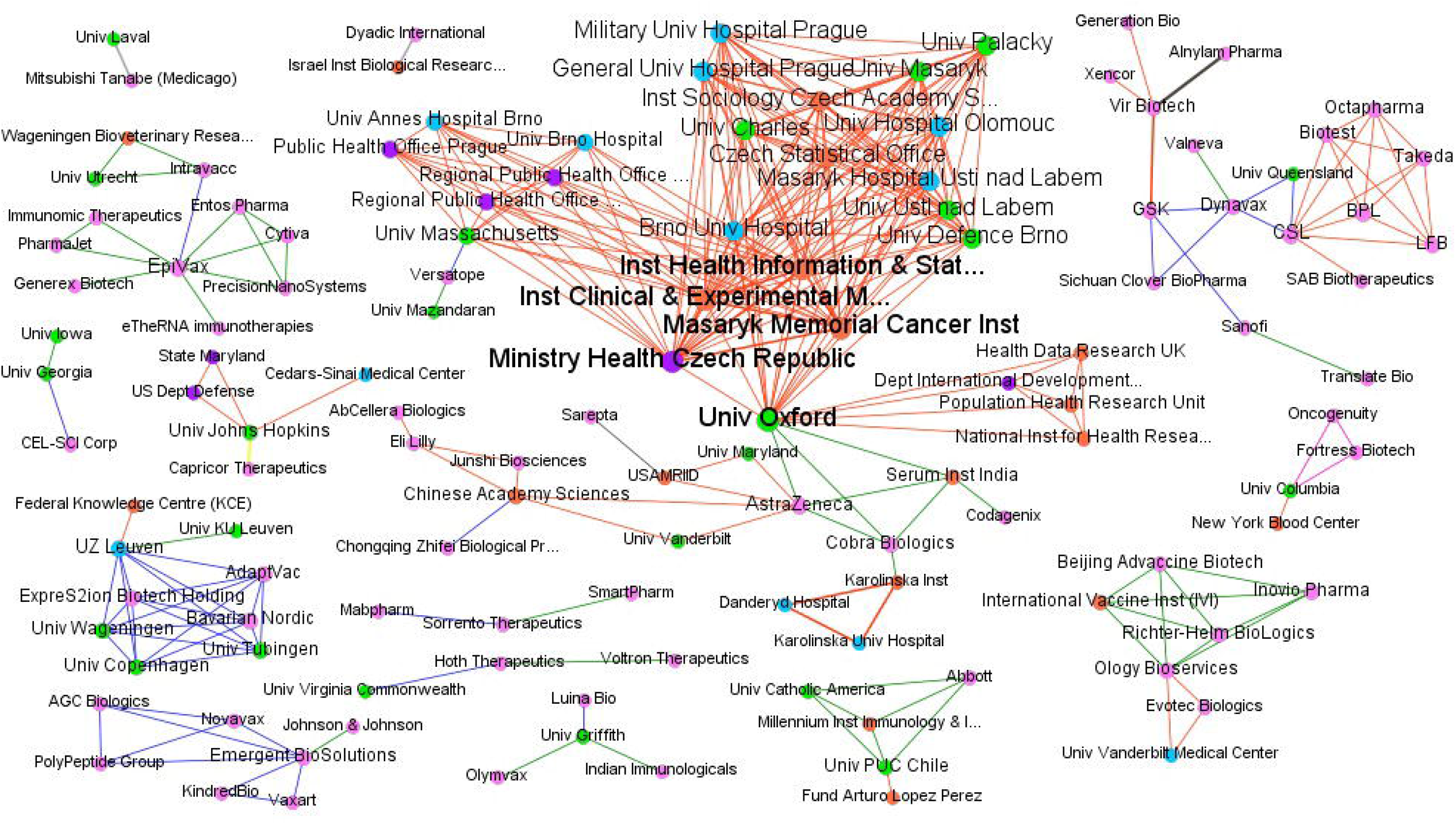
CT cooperation on Antibody CT. Legend: Pink = Firm; Blue = Hospital; Green = University; Orange = Research Institute; Red = Government

Antibodies were the only category that brought together the 12 government organizations in the network. For this category, Vir Biotech, and the Instituto Nacional de Enfermedades Respiratorias (INER) stand out, with partnerships in four and three CTs, respectively. While the INER conducts research with hospitals and other RIs on convalescent plasma, Vir Biotech works in conjunction with three other firms on neutralizing antibodies. It even uses its monoclonal antibody platform in two partnerships with GSK, which has experience in functional genomics.

The second largest cluster is vaccines (Fig 6), formed mainly by CTs between university, firms, and RIs, mainly focused on recombinant DNA, RNA, and live attenuated virus. EpiVax and Tonix Pharma stand out with four partnerships each. Tonix maintains cooperation with the Southern Research Institute and has three other relationships with the University of Alberta, all involving engineered live attenuated virus in the preclinical phase. In turn, EpiVax cooperates with seven organizations, each with a different expertise.

**Fig 6.**
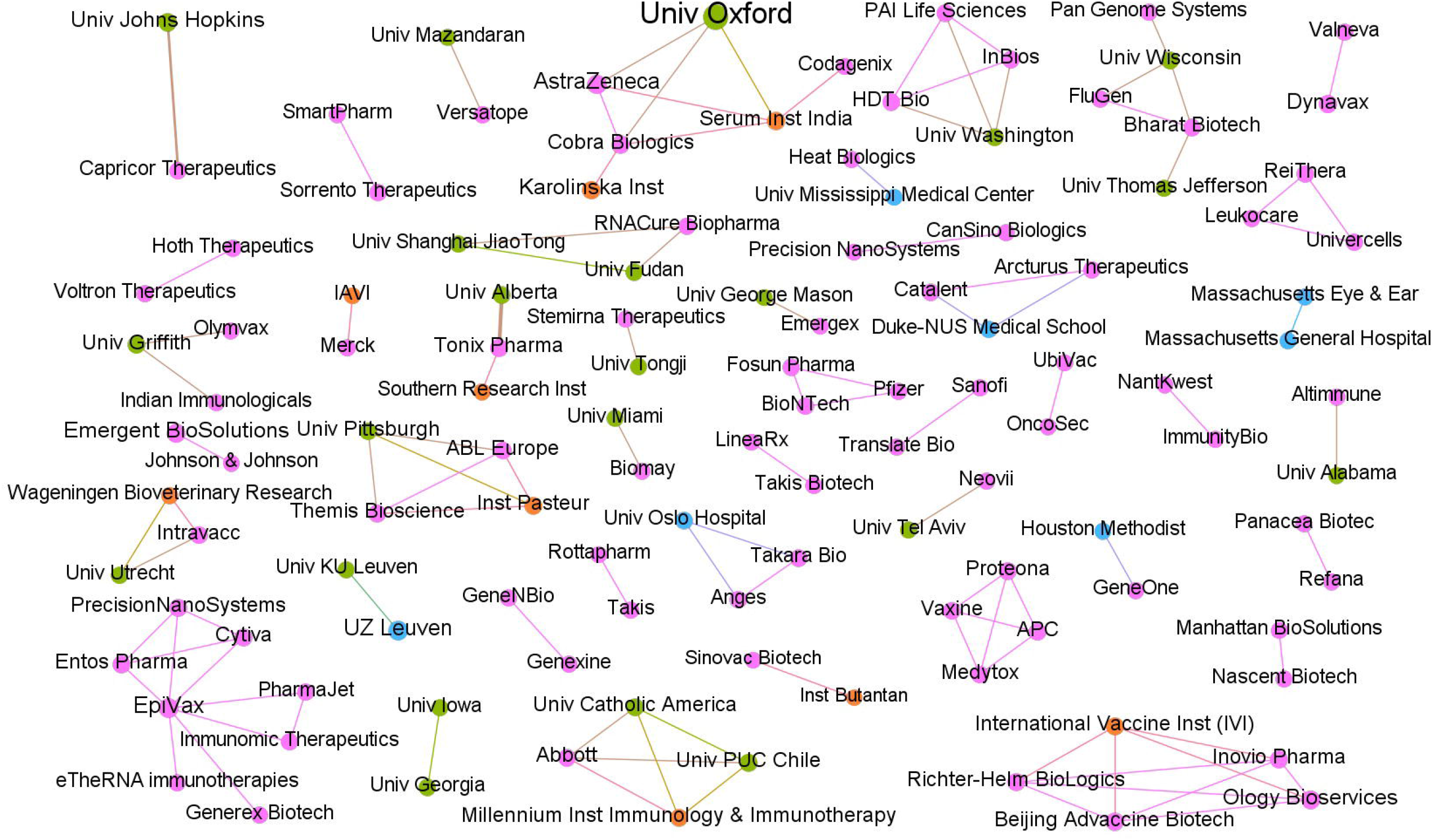
CT cooperation on Vaccine.

Among the more advanced trials, four vaccines stand out. The AZD1222 vaccine candidate, based on an adenoviral vector, is in a phase III CT through a partnership between the University of Oxford and AstraZeneca, with the participation of Cobra Biologics and Serum Institute. The mRNA-1273 (phase III), developed by Moderna with support from the NIH, uses messenger RNA technology for the expression of the S protein of SARS-CoV-2. Also, in phase III CT is an inactivated SARS-CoV-2 vaccine candidate developed by in a partnership with the Butantan Institute in Brazil. The partnership between Pfizer and BioNTech also generates a product based on messenger RNA, which is being tested in a phase II CT in Germany.

There are 30 protein-based technologies (Fig 7) which can also include protein subunit vaccines, of which 28 are in the preclinical phase and only two are in phase I CTs. The CT involving SCB-2019, which is also a protein subunit vaccine by Sichuan Clover BioPharma combines the expertise of GSK and Dynavax to increase the immune response of patients. The NVX-CoV2373, a vaccine developed by Novavax is a candidate which include partnerships with PolyPeptide Group, AGC Biologics and Emergent BioSolution to ensure the scaling of compounds necessary for final product development and large-scale manufacturing. These trials show that firms have intensified cooperation relationships to foster a faster response to COVID-19 and ensure product scalability.

**Fig 7.**
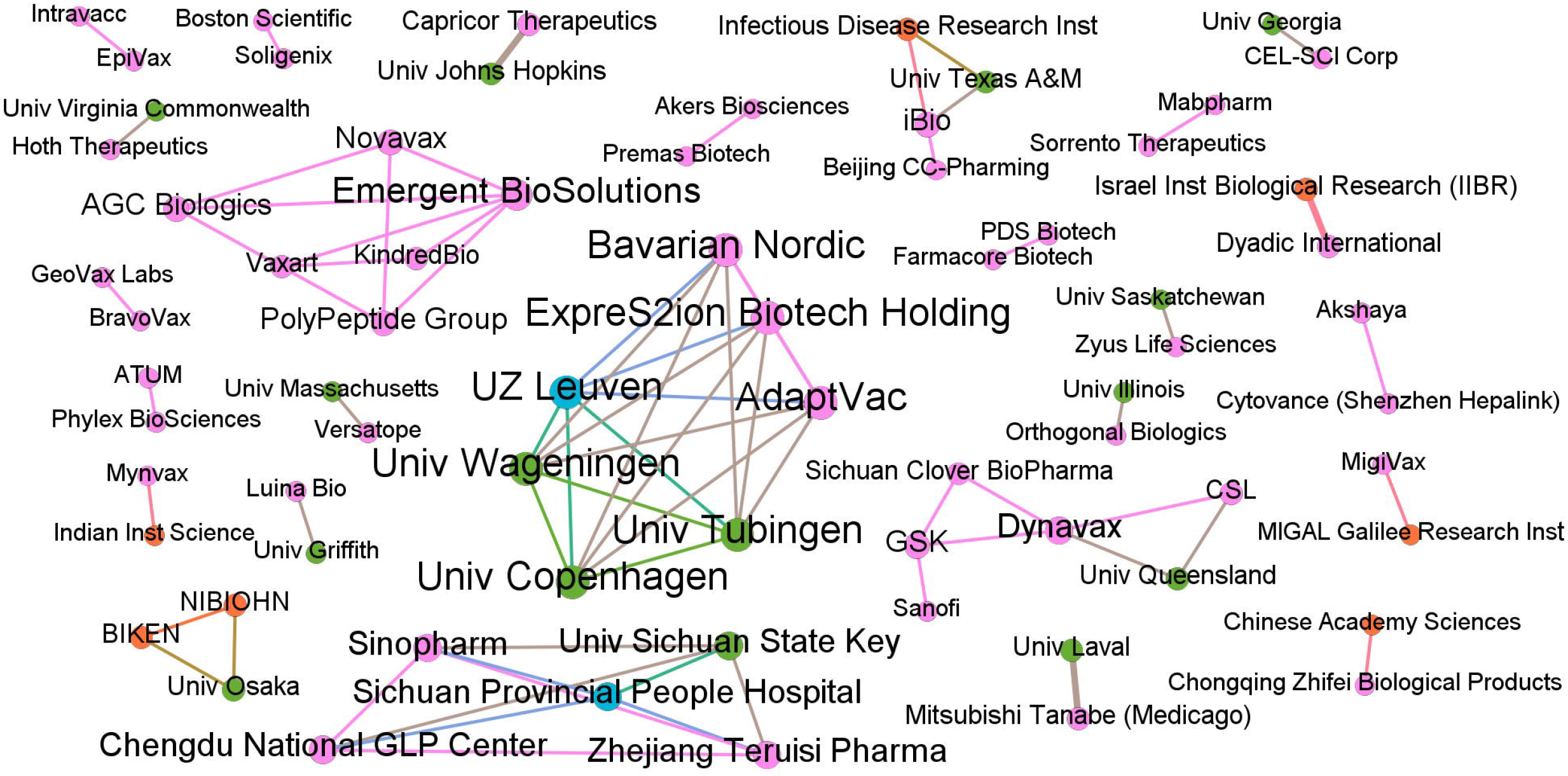
CT cooperation on Protein-based.

The CT of siRNA, cell therapy, small molecule, and nucleic acid (Fig 8) represent approximately 11% of the partnerships. Among the six CTs focused on siRNA, three stand out for the relationship between Alnylam Pharma and Vir Biotech and the combined use of lung delivery of novel conjugates of siRNA and expertise in infectious diseases. BioMed, Mannin Research, and Cyclica focus on small immunosuppressants molecules to reduce the effects of the symptoms caused by the SARS-COV-2 infection.

**Fig 8.**
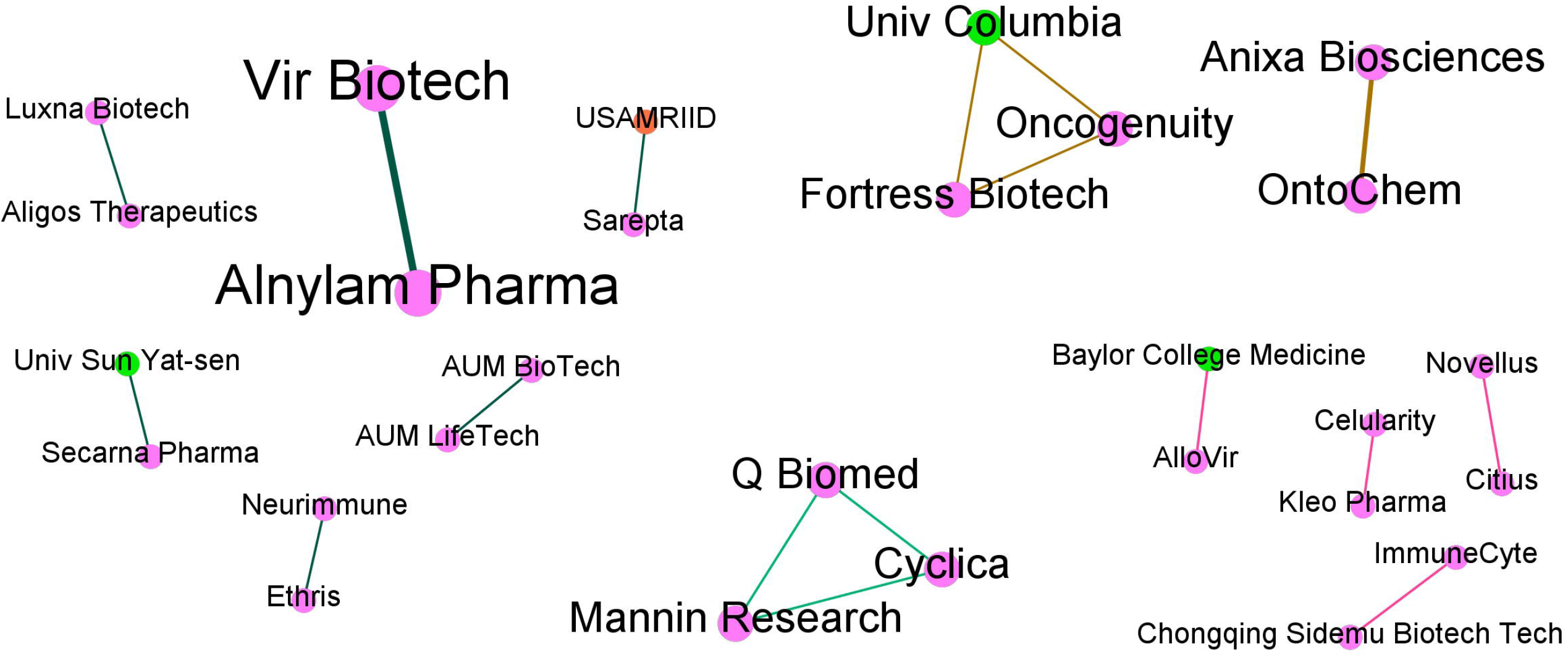
CT cooperation on iRNA/ Cell therapy/ Small mollecule.

Among the four CTs involving cell therapy, the Chongging Biotechnology and ImmunCyte (phase I/II) partnership stands out for CAR-modified NK cells, which recognize and eliminate the virus. Intraregional partnerships in China, in the biomedical field, favour technology spillover abroad and the production of innovation [27].

Four firms and one university developed three preclinical trials based on nucleic acids. OntoChem and Anixa Biosciences participate in two trials involving technology that inhibits the ability of the virus to replicate and bind to human cell proteins. Conversely, the University of Columbia, Oncogenuity, and Fortress Biotech use a platform to produce oligomers that can help fight COVID-19 and accelerate the discovery of treatments for new outbreaks.

The intense participation of Chinese organizations in COVID-19 CTs is notable, which may represent not only a change in the geographical axis of the development of this type of technology but also the emergence of new players in the fierce biopharmaceutical market. The Chinese firms had also developing the therapeutic strategies before any other country, leading the landscape since the outbreak starts overthere. Furthermore, organizations from countries without rich traditions in this area, based in Eastern Europe and Latin America, have started to participate in this type of development.

Some organizations operate in more than one category (Fig 9), which may be an indication of greater capacity for technological development and partnerships. The choice to cooperate in different fields, with different partners, contributes to expanding the connection of CT development networks for COVID-19, directly interfering in the network metrics, which will imply more relevant nodes.

**Fig 9.**
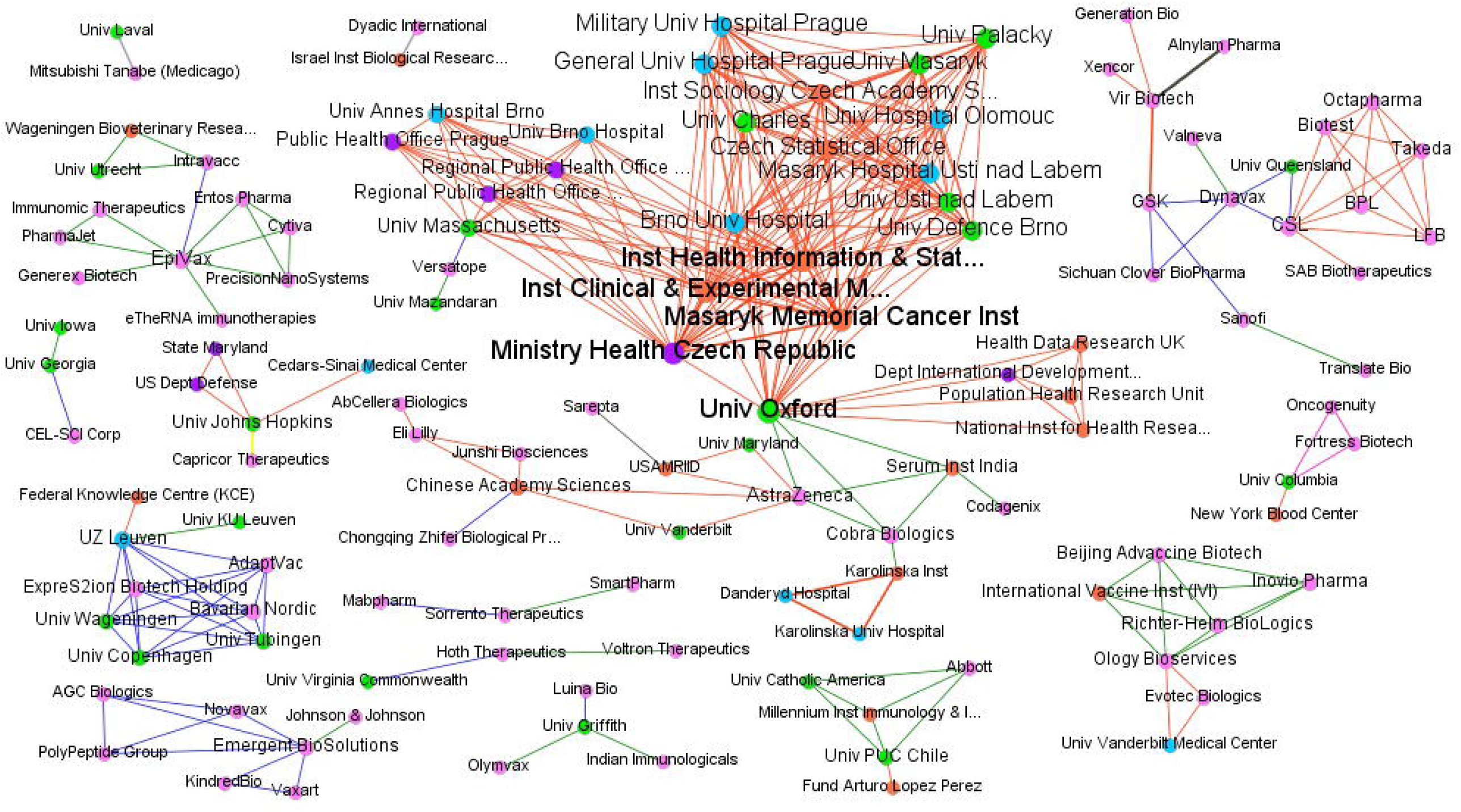
Cooperation network for the development of more than one category of CT.

The network metrics (Table SM 3) indicate a low density, with an average and weighted degree with similar values, because only 64 institutions developed at least two CTs. Among these, only 37 institutions have built partnerships with different players. The density of the less explored categories can bring an understanding that there are more connections, but this occurs due to the presence of few actors, requiring fewer connections [28]. The diameter of the overall network is eight connections, which is considered a high value, impacted by organizations that have different partners in more than one CT.

Regarding betweenness centrality, which indicates high influence within a network due to the power to mediate relationships, it was found that 38 organizations influence connectivity, including 18 firms, nine universities and eight RIs (Table SM 4). Among these organizations, ten developed partnerships in only one category, as they chose to focus their efforts on projects with the same technological direction.

UZ Leuven and Johns Hopkins University cooperate in three different categories and complement their expertise as well as diversify their efforts. The University Hospital of Leuven established three partnerships, one with the University of Leuven, which is focused on a vaccine, another with the Belgian Federal Knowledge Centre, which is focussed on an antibody, and another involving six actors, which involves testing the protein developed by ExpreS2ion Biotech. Of the four CTs in which Johns Hopkins University cooperates, two involve partnerships with Capricor Therapeutics, with aims of advancing protein-based therapies and vaccines and both with a focus on nanoparticles. This university also has prominent partnership with the State of Maryland and the US Department of Defence.

The remaining five organizations with the highest betweenness centrality are interconnected. The University of Oxford has the highest betweenness centrality, as it participates in a CT that brings together 18 other organizations, in addition to acting in two other trials with partners that also positively influence the network connection, e.g., AstraZeneca. This firm also participates in three CTs and sought to establish more specific partnerships with organizations that have more than one CT in cooperation. The high volume of triangulations of some organizations suggests multiple partnerships with a high impact on centrality and low risk diversification.

When analysing the strength of cooperation, of the 700 relationships, 391 (56%) are one-to-one relationships; that is, organizations participating in only one CT and with a single partner. Therefore, for these cases, SC and UDC have the maximum value (1). Furthermore, 94% of the partners have at least one party with a high dependence on the relationship (UDC in Q4). These characteristics of a high degree of cooperation and asymmetry in unilateral dependence can be attributed to partnership models with well-defined specializations between the parties, which occur in extremely competitive environments, corresponding to the current moment of the pandemic.

For the other partners (309), SC was found in Q1 only for one relationship, i.e. AstraZeneca and Chinese Academy Sciences, that are developing a neutralizing antibody, referring to the fact that both work independently of each other. The other organizations have a median cooperation affinity, with a tendency to increase. This influences the distribution of quartiles of the UDC, whose predominance in the combinations Q1-Q4 and Q2-Q4 reinforces the dependence asymmetry in the cooperation relationships. This finding is evident when observing the SC and UDC measurements by category and type of organization, where antibodies, vaccines and proteins fit this profile (S5-6 Table).

Among the organizations with greater cooperation in CTs, Vir Biotech stands out with seven trials in progress, of which three focus on siRNA and four focus on antibodies. Vir Biotech showed greater cooperation affinity with GSK for antibodies (SC = 0.44) and Alnylam Pharma for siRNA (SC = 0.65). Despite this greater affinity, Vir Biotech has a low dependence on the relationship, unlike the other two partners, who have a high dependence. Johns Hopkins University, with four trials involving antibodies, one involving proteins, and one involving a vaccine, shows a greater cooperation affinity with Capricor (SC = 0.58) and has a low UDC (0.33), while its partner has a relationship of total dependence in this category. Another organization that stands out in this analysis is EpiVax, with six ongoing trials, of which five involve vaccines and one involves a protein, with low affinity with its partners but with dependence on the cooperation relationship.

## Discussion

Understanding the cooperation dynamics of organizations allowed us to assess of how crisis situations like this pandemic can influenciated the structure of cooperation networks. Thus, it was possible to understand the characteristics of these networks, how they are established, which are the partnerships profiles and features and the types of technologies and research developed.

The findings pointed out that both general and cooperation amount of CT are concentrated on antibody, vaccine, and protein categories. The Oxford University, Astrazeneca, EpiVax, Vir Biotech, University of Johns Hopkins and Chinese Academy of Science stand out for acting strongly in cooperation with other organizations. It is worth highlighting the advent of Chinese organizations in this sector, with intense cooperation. This indicates the emergence of new players in the biotechnology field and, even, a possible change in the geographical axis of this type of technology development.

There is a high collaboration rate in the CT networks, but it does not reflect to a complete open innovation practice, since the organizations have specific partnerships, that restrict a wide cooperation network with more flow of information and knowledge. However, it can be said that the challenge of finding therapeutic or immunological solutions for COVID-19 demonstrates a clear composition of cooperation groups that complement their skills to manage their interests and organizational strategies to beat the pandemic. Thus, there are predominantly vertical cooperation networks in which organizations are from the same productive chain. It is noted that the geographical barriers, although still strong, lose relevance in this new context when the cooperation of organizations from different countries is established, making possible the emergence of technological development in countries with soft tradition in this field, such as some based in Eastern Europe and Latin America. In addition, there is a high strength of cooperation between organizations in CT and an asymmetry in the unilateral dependence on partnerships.

Another characteristic observed is the wide diversity of partnerships between firms, hospitals, RI, and universities. The role of universities has a special meaning, which in the context of a pandemic requires complex research and leads to intensification of partnerships for CT execution, being a complementary way to achieve the desired objetives. Hospitals are also seen as key organizations in this type of partnership in order to enable, quickly and in multiple centers facilitating the testing of new drugs and vaccines. Finally, in the case of firms, they set aside their history of disputes and started to share competences. In this new paradigm, there can be partnerships not only in CT but also in pre-competitive technologies development. This experience is expected to change the way of organizations define their R&D strategies and start to adopt more widely a collaborative innovation model.

This study has some limitations due to the availability of information. There is a difference between the quality of information provided by universities and industries. These variations can be perceived according to the geographical locations of organizations and clinical studies. In addition, there is the possibility that some groups may not fully report their status for competitive reasons or overreport to attract more funding. In future research, it is intended to analyze more broadly the process of technological development for drugs and vaccines that will result in disruptive technologies protected by patents, complementing the understanding of different aspects of cooperation in this new context such as the current pandemic.

## Supporting Information

**S1 Appendix. Main constructs used for innovation, SNA, and clinical trials**.

**S2 Appendix. Strength of Cooperation (SC)**

**S1 Table. Grouping of categories to characterize clinical studies**. The categories were grouped considering the technological proximity that exists between them. In the case of vaccines, the class information and the type of compound provided by the Biocentury were also evaluated.

**S2 Table. Characterization of studies by category**.

**S3 Table. Metrics of the general network and grouped category**.

**S4 Table. Metrics regarding the relevance of the nodes**.

**S5 Table. General statistics of strength of cooperation**.

**S6 Table. Analysis of the strength of cooperation and unilateral dependence by category and type of organization**

